# Application of *hsp60* amplicon sequencing to characterize microbial communities associated with juvenile and adult *Euprymna scolopes* squid

**DOI:** 10.1101/2024.09.23.614625

**Authors:** Steph Smith, Clotilde Bongrand, Susannah Lawhorn, Edward G. Ruby, Alecia N. Septer

**Author notes:** **Competing Interests** The authors declare no competing interests.

## Abstract

The symbiotic relationship between *Vibrio (Aliivibrio) fischeri* and the Hawaiian bobtail squid, *Euprymna scolopes*, serves as a key model for understanding host-microbe interactions. Traditional culture-based methods have primarily isolated *V. fischeri* from the light organs of wild-caught squid, yet culture-independent analyses of this symbiotic microbiome remain limited. This study aims to enhance species-level resolution of bacterial communities associated with *E. scolopes* using *hsp60* amplicon sequencing. We validated our *hsp60* sequencing approach using pure cultures and mixed bacterial populations, demonstrating its ability to distinguish *V. fischeri* from other closely-related vibrios and the possibility of using this approach for strain-level diversity with further optimization. This approach was applied to whole-animal juvenile squid exposed to either seawater or a clonal *V. fischeri* inoculum, as well as ventate samples and light organ cores from wild-caught adults. *V. fischeri* accounted for the majority of the identifiable taxa for whole-animal juvenile samples and comprised 94%-99% of amplicon sequence variants (ASVs) for adult light organ core samples, confirming that *V. fischeri* is the dominant, if not sole, symbiont typically associated with *E. scolopes* light organs. In one ventate sample, *V. fischeri* comprised 82% of reads, indicating the potential for non-invasive community assessments using this approach. Analysis of non-*V. fischeri* ASVs revealed that *Bradyrhizobium spp*. and other members of the Rhodobacterales order are conserved across juvenile and adult samples. These findings provide insight into the presence of additional microbial associations with the squid host tissue outside of the light organ that have not been previously detected through traditional culture methods.

## Brief Communication

Symbiosis model systems have been essential to expand our understanding of how microbes interact with their hosts. The symbiosis between *Vibrio fischeri* and *Euprymna scolopes* (Hawaiian bobtail squid) is a model for studying host-bacterial interactions in the context of a beneficial association. Juvenile *E. scolopes* hatch without their bacterial symbiont, *V. fischeri*, which they acquire from the surrounding seawater. Independent *V. fischeri* strains quickly colonize the crypt spaces within the bi-lobed symbiotic light organ where their bioluminescence aids in the squid’s nocturnal activity [1].

Although *V. fischeri* is the only species isolated from the light organs of wild-caught animals, only one study has applied a culture-independent approach for examining the diversity of bacteria associated with adult animals. This study, which applied amplicon sequencing of the V3-V4 region of the 16S *rRNA* gene to the light organs of two wild-caught adult squid, found the majority of reads (79%) were most similar to *V. fischeri*, with another 5% most similar to *Vibrio litoralis* [2]. Although this study was the first to apply a culture-independent approach to the light organ symbiosis, it also revealed a need for using an amplicon target that better distinguishes between *Vibrio* species, to increase confidence for assigning species-, and possibly strain-level taxonomy to amplicon sequence variants (ASVs).

The goal of this work was to develop a protocol to apply *hsp60* amplicon sequencing to the vibrio-squid symbiosis. The *hsp60* gene encodes a conserved protein that serves as a better marker for distinguishing and identifying species-level classification within the closely-related Vibrionaceae [3, 4]. We applied this approach to pure bacterial cultures, as well as whole-animal juvenile *E. scolopes* squid, juvenile ventate water, and light organ cores from four wild-caught adults. Based on these results, we propose how this approach can be applied to the vibrio-squid system, its limitations, and how further optimization could expand its utility.

We first applied *hsp60* amplicon sequencing to known culture samples. DNA extraction and amplicon sequencing was performed on three types of culture samples: 1) a single strain of *V. fischeri*, 2) mixed cultures of multiple *V. fischeri* strains, and 3) a multispecies culture, to validate that *V. fischeri* can be distinguished from other species and observe how strain-level variance might appear in the amplicon sequencing analysis. Because this approach involves amplification of the target gDNA sequence, amplification or sequencing errors could introduce nucleotide changes and thus result in distinct ASVs from a single, clonal copy of the target, or natural within-population variation of the target sequence could result in multiple ASVs, despite representing a single bacterial lineage [5-7].

Analysis of the culture samples indicate that *V. fischeri* ASVs can be distinguished from ASVs for other species (Fig 1A), while strain-level resolution was less clear (Fig 1B). Although most *V. fischeri* genomes encode a unique *hsp60* sequence (Fig 1C), including the strains used in our culture samples (ES114, MB13B1, and MB13B2), the *hsp60* amplicon sequencing results yielded more than one unique ASV per input strain. At the species level, all 18 unique ASVs were assigned to *V. fischeri*. However, at the strain level, six unique ASVs were identified as ES114, 11 ASVs were identified as MB13B2, and one ASV was identified as MB13B1. When the dataset was filtered to exclude ASVs that represent less than 0.1% of total reads across any given sample, two dominant unique ASVs were identified for both ES114 and MB13B2, and a single unique ASV was identified for MB13B1 (Fig 1B), suggesting that removal of low-abundance ASVs that may result from sequencing errors does not resolve the mismatch between the number of strains (three) and the number of unique ASVs (five) for the known culture samples.

**Figure 1.**
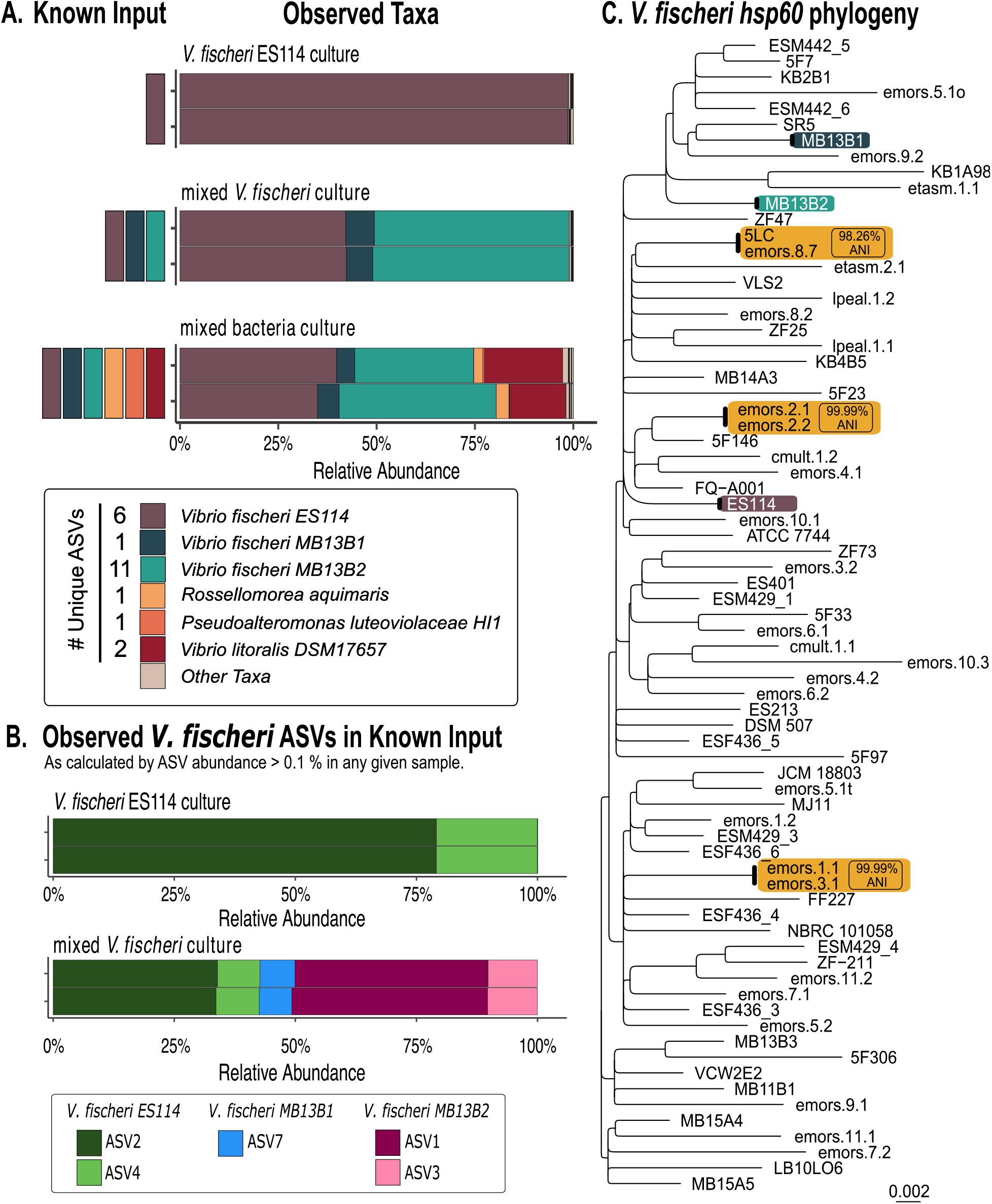
*hsp60* amplicon sequencing differentiates *Vibrio fischeri* from other closely related vibrios at the species level. **(A)** Known Input corresponds to bacterial isolates added to each culture sample, and Observed Taxa reports the relative abundance (%) of taxa identified from each cultured sample using *hsp60* amplicon sequencing. All known input strains were detected in their corresponding samples, including in mixed cultures comprised of multiple *Vibrio fischeri* strains and closely related vibrios. The number of unique ASVs that were assigned to each taxonomic identifier is reported here. Stacked bars indicate technical replicates of each treatment. **(B)** Relative abundance (%) of *V. fischeri* ASVs comprising > 0.1% of total abundance in any given sample. Of ASVs that meet this criterion, two ASVs were identified as *V. fischeri* ES114, two ASVs were identified as *V. fischeri* MB13B2, and one ASV was identified as *V. fischeri* MB13B1. Abundance of any additional unique ASVs identified in panel A fell below the abundance cutoff of 0.1%. **(C)** Neighbor-joining phylogenetic tree analysis based on distance matrix calculated by multiple sequence alignment (MSA) of 73 *V. fischeri hsp60* sequences obtained from NCBI and trimmed to the region amplified by the primers used in this study (see Table S1). Orange boxes represent strains with 100% identity between *hsp60* sequences compared to the average nucleotide identity (ANI) between the corresponding strains at the whole-genome level as calculated by FastANI (98.26 – 99.99% ANI).

We conclude that, although *hsp60* amplicon sequencing can distinguish *V. fischeri* from other bacterial species, further work is needed to identify appropriate computational thresholds to separate true strain-level sequence variants from ASVs that may arise from errors in amplification or sequencing. Since we are currently unable to determine whether minority ASVs represent unique strains or artifacts of the approach, we chose not to apply strain-level diversity analyses to the animal samples in the following section.

We next applied *hsp60* amplicon sequencing to animal samples. DNA extraction and amplicon sequencing was performed on three types of samples: 1) homogenized whole animal juveniles colonized with Hawaiian seawater or a single *V. fischeri* strain, 2) juvenile ventate water, or 3) paired cores of light organ lobes (A and B) from four wild-caught adult squid. Although only 31% and 62% of the ASVs associated with juvenile squid were identified as *V. fischeri*, the remaining ASVs had no assigned taxonomy, making *V*.

*fischeri* the majority of identified reads in these samples. The ventate sample results were variable: one sample returned 82% of reads identified as *V. fischeri* while the second ventate sample was comparable to juvenile squid that were not exposed to *V. fischeri* (aposymbiotic; apo) (Fig 2A). These results suggest this approach could be optimized by plating a subsample of the ventate prior to extraction to verify the animals did in fact vent. For adult samples, *V. fischeri* ASVs comprised 94% -99% of the total reads, confirming that *V. fischeri* is the primary symbiont in the light organ using a culture-independent technique (Fig 2A).

**Figure 2.**
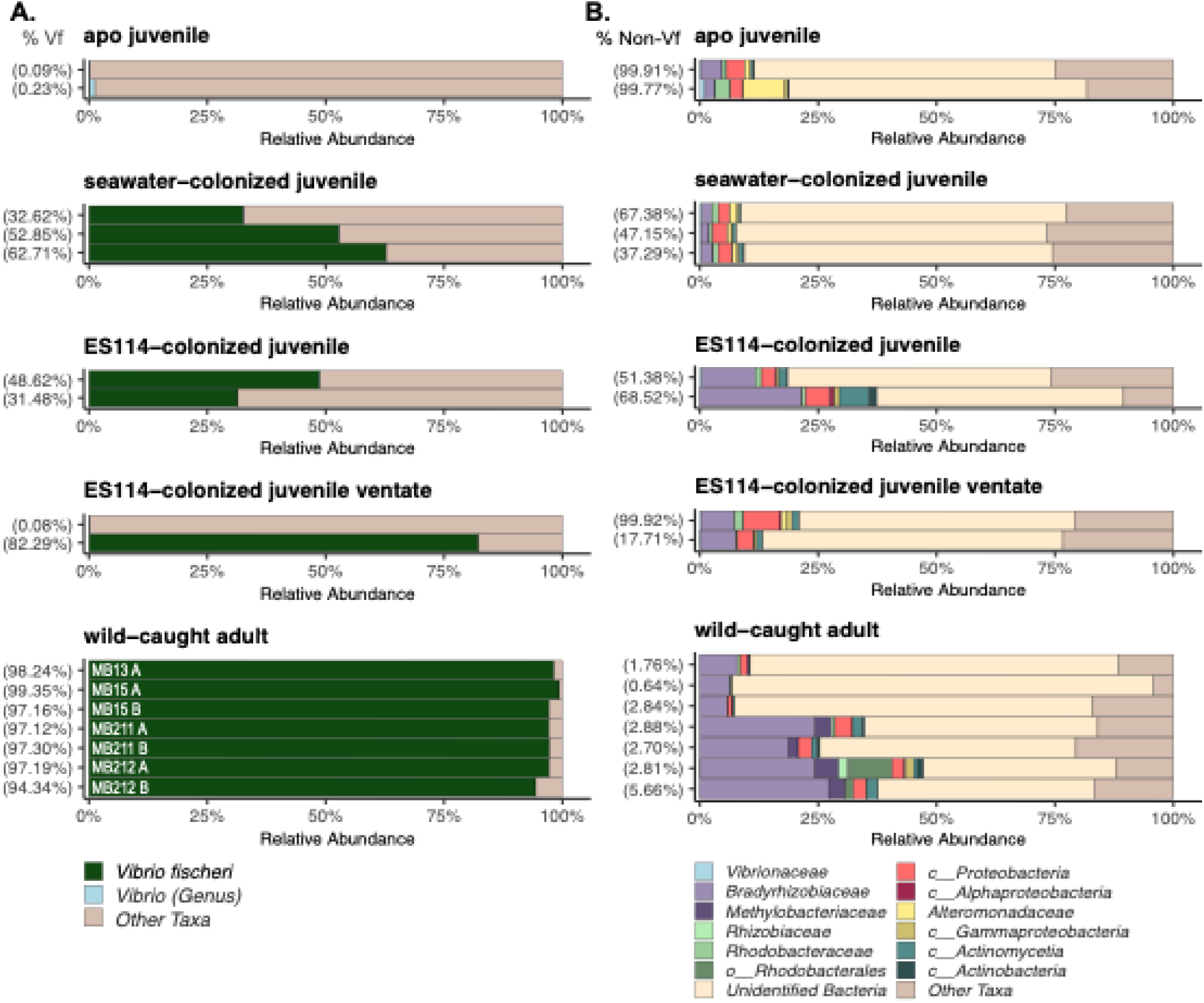
Relative abundance of *V. fischeri* and other conserved taxa associated with juvenile and adult squid. **(A)** Relative abundance of *V. fischeri*, other vibrios, and non-vibrio taxa in juvenile squid, juvenile squid ventate, and adult squid. The percentage of reads identified as *V. fischeri* in each sample is indicated on the y-axis (% Vf). In wild-caught adult panel, MB* indicates animal identifier, with A or B indicating paired samples from each lobe of the same light organ. **(B)** Corresponding relative abundance of non-*V. fischeri* taxa in the samples described in panel A. Taxa are plotted at the family level, and *Vibrionaceae* excludes ASVs identified as *V. fischeri*. Unidentified Bacteria correspond to ASVs that were not assigned taxonomy past the kingdom level, likely due to a lack of similar sequences in available *hsp60* reference sequences. Other Taxa correspond to ASVs that were assigned taxonomy to at least the family level but make up < 0.1 % of the total read abundance. Sample order is consistent between panel A and panel B.

Finally, we explored the ASVs that were not identified as *V. fischeri* in animal samples, which comprised 37-68% of reads for juveniles and 0.6-5% of reads for adult light organ cores. This result was not unexpected given that these samples consist of either whole animal homogenates (juvenile) or light organ cores (adult), and any bacteria associated with the surface of the light organ or other tissue would be amplified along with the light organ population. Analysis of these non-*V. fischeri* ASVs revealed two important findings: 1) the majority of these ASVs were not assigned a taxonomy (likely due to lack of similar sequences available in *hsp60* databases), and 2) several conserved taxa appeared in both juvenile and adult animals that were dominated by members of the Rhodobacterales order, including *Bradyrhizobium spp*. (Fig 2B). Importantly, none of these conserved ASVs were detected in either of our negative extraction control (NEC) samples, and therefore are unlikely to represent contaminants. These conserved taxa may represent another important microbial association outside of the primary light organ symbiosis. *Bradyrhizobium* have primarily been studied as plant symbionts known for their ability to fix nitrogen, with a recent isolate encoding genes to assimilate ammonium [8], which is also excreted by the squid host [9, 10]. Future work may permit the cultivation of such taxa and the exploration of its potential localization and interactions with the squid host.

## Methods

See supplemental methods for details pertaining to squid collection and experimentation, bacterial strains, DNA extraction, sequencing, and bioinformatic analyses.

## Acknowledgements

This work was supported by NIGMS grants R35 GM137886 awarded to ANS and R01 GM135254 to EGR. SNS was supported by the National Science Defense and Engineering Graduate fellowship (NDSEG) and the UNC EMES Mills Brown Fellowship Award. We thank the UNC High Throughput Sequencing Facility (HTSF) for their support and guidance in this work.

## Data Availability

Amplicon sequences are available at NCBI under BioProject PRJNA1136500. (Reviewer link here: https://dataview.ncbi.nlm.nih.gov/object/PRJNA1136500?reviewer=qqgcnl6oe4dkt3djps1p735421).

## Competing Interests

The authors declare no competing interests.

## Notes

### Competing Interest Statement

The authors have declared no competing interest.

