## Supplemental Materials for "Application of *hsp60* amplicon sequencing to characterize microbial communities associated with juvenile and adult *Euprymna scolopes* squid"

### Supplemental Methods

#### Adult squid collection.

Adult *Euprymna scolopes* light organ cores were previously obtained by collecting wild-caught animals from Maunalua Bay with a dipnet, euthanized, ventrally dissected, and cores from each light organ lobe taken and stored at -80°C until DNA extraction.

#### Juvenile squid colonization.

Juvenile *Euprymna scolopes* were hatched into bacteria-free seawater and exposed to either a clonal inoculum of *V. fischeri* or to natural seawater. Animals were euthanized and whole animals were homogenized and stored at -80°C prior to DNA extraction.

#### Bacterial strains.

Bacterial isolates used in this study include *V. fischeri* ES114, *V. fischeri* MB13B1, *V. fischeri* MB13B2, *Rossellomorea aquimaris* TF-12, *Pseudoalteromonas luteoviolacea* HI1, and *Vibrio litoralis* DSM17657. Individual overnight cultures were grown at 28°C in Luria-Bertani salt (LBS)[1] media and diluted to 1.0 $OD_{600}$.

In mixed culture treatments, equal volumes of individual cultures were combined to a total volume of 1 mL. Clonal treatments are comprised of 1 mL of the respective normalized culture. Individual replicates of each culture treatment were pelleted using centrifugation and stored at -80°C until DNA extraction.

For juvenile squid colonization experiments, individual overnight cultures were diluted 100x in seawater tryptone (SWT)[1] medium broth and grown with shaking at 28°C until mid-exponential phase (0.5 $OD_{600}$). Bacteria were diluted to a concentration of ~5,000 cells/mL in 50 mL of filter-sterilized ocean water (FSOW) in which newly hatched juvenile squid were transferred. Incubations were performed at room temperature, between 21 and 24°C.

#### DNA extraction and *hsp60* amplicon sequencing.

DNA was extracted simultaneously from all samples using a Zymo Quick-DNA Fungal/Bacterial Miniprep kit (Zymo Research, Irvine, CA, USA) along with duplicate negative extraction controls. Amplicon sequencing of the *hsp60* gene was performed by UNC's High Throughput Sequencing Facility on a MiSeq System using primer sets H279/H280 and H1612/H1613 mixed at a 1:3 molar ratio as previously described [2-4]. See table S2 for amplicon read sequences processing summary.

Demultiplexed FastQ files were processed with DADA2 v1.10 [5]. The following parameters were used for sequence filtering and trimming: trimLeft=26, maxN=0, maxEE=2. The parameter trimLeft=26 was used to uniformly remove the primer sequence from demultiplexed forward reads. Due to amplicon length, only forward reads were analyzed in this study. Chimeric sequences were removed with the removeBimeraDenovo function of DADA2. Taxonomic classification of each ASV was performed with a naïve Bayesian classifier in QIIME2 [6] using p-confidence=0.6 and a custom *cpn60* reference database, available upon request (cpn60_classifier_v11.qza). This reference database comprises publicly available *cpn60* reference sequences [7] appended with the *hsp60* sequence and taxonomy of each bacterial isolate used in this study (*V. fischeri* ES114[8], *V. fischeri* MB13B1[9], *V. fischeri* MB13B2[9], *Rossellomorea aquimaris* TF-12[10], *Pseudoalteromonas luteoviolacea* HI1[11], and *Vibrio litoralis* DSM17657[12]). The identification of sequencing contaminants was assessed using decontam v1.12 [13] based on frequency and input DNA concentration, as well as prevalence in negative extraction controls. Diversity analyses were performed with PhyloSeq v1.40.0 [14].

Relative abundance was assessed with the microViz v0.12.4 package [15]. See Escolopes_hsp60_Analysis for Rmd documentation of the bioinformatic analyses performed.

**Data availability statement.**
The amplicon sequences presented in this study are available via NCBI under BioProject PRJNA1136500.
Reviewer link:
https://dataview.ncbi.nlm.nih.gov/object/PRJNA1136500?reviewer=qqgcnl6oe4dkt3djps1p735421

**Ethics statement.**
The University of Hawaii Institutional and Animal Care and Use Committee (IACUC) is only allowed to review research using vertebrate animals. Author ER has a letter from the University veterinarian that states the use of cephalopods in the research conducted in this study would pass IACUC standards if they were allowed to formally review it.

**Supplemental References.**
